# When to attend? Temporal attention interacts with expectation

**DOI:** 10.1101/2023.12.08.570819

**Authors:** Aysun Duyar, Shiyang Ren, Marisa Carrasco

## Abstract

Temporal attention is voluntarily deployed at specific moments, whereas temporal expectation is deployed according to timing probabilities. When the target appears at an expected moment in a sequence, temporal attention improves performance at the attended moments, but the timing and the precision of the attentional window remain unknown. Here we independently and concurrently manipulated temporal attention–via behavioral relevance–and temporal expectation–via session-wise precision and trial-wise hazard rate–to investigate whether and how these mechanisms interact to improve perception. Our results reveal that temporal attention interacts with temporal expectation–the higher the precision, the stronger the attention benefit, but surprisingly this benefit decreased with delayed onset despite the increasing probability of stimulus appearance. When attention was suboptimally deployed to earlier than expected moments, it could not be reoriented to a later time point. These findings provide evidence that temporal attention and temporal expectation are different mechanisms, and highlight their interplay in optimizing visual performance.

**Relevance:** Our ability to process visual information is limited both across space and time. Here we disentangle how two mechanisms–attention and expectation–help us overcome temporal limitations. We concurrently manipulated attention and expectation independently to investigate whether and how they interact. We found that temporal attention interacts with two distinct forms of expectation. Temporal expectation strengthens the benefits of temporal attention on performance for the attended time points, depending on how precise the expectations are. Surprisingly, the advantages of attention decrease when stimuli occur later than expected, suggesting a limitation of attention to reorient from earlier to later time points. This study provides further evidence that humans cannot sustain temporal attention even over short periods, reveals that although temporal attention and expectation interact to improve visual performance, expectation suboptimally guides attention, and highlights that attention and expectation are different temporal mechanisms.

## INTRODUCTION

The visual environment is constantly changing, and our brain receives more sensory information than it can process over time due to its limited temporal capacity (Shapiro, 2001). Temporal attention and expectation are two key cognitive mechanisms that provide shortcuts to optimize perception.Endogenous temporal attention is the voluntary prioritization of task relevant time, whereas temporal expectation refers to the probability of event timing (e.g., de Lange et al., 2018; Denison et al., 2021; Duyar et al., 2023; Palmieri et al., 2023).

Temporal expectations, which are based on the predictability of event timing, increase visual accuracy and decrease reaction times (Rohenkohl et al., 2012; Vangkilde, et al., 2012). Expectations are developed through the statistical regularities of external events embedded within noisy distributions, resulting in uncertainty regarding event timing. Uncertainty, as a mathematical term, is measured by the variance of a probability distribution (Feldman & Friston, 2010). Temporal uncertainty, which is inversely related to the precision of the expectation at a given moment, impairs the performance in visual and auditory modalities (Abeles et al., 2020; Egan et al., 1961; Rolke & Hofmann, 2007). Another source of temporal expectation is hazard rate: The increasing probability over time of an event occurring, given that it has not yet occurred within a certain time window, facilitates online updating of expectation (Duyar et al., 2023; Grabenhorst, et al., 2021; Nobre et al., 2007).

Temporal attention enables us to select and prioritize specific moments when relevant events occur and enhances the processing of the sensory information at such moments (Coull, 2004; Denison, 2023; Griffin et al., 2001; Nobre, 2001, 2007). These enhancements in visual perception, which go beyond mere expectation, result in impairments at the unattended moments (Denison et al., 2017, 2021). Hence, temporal attention allocation results in tradeoffs–benefits at the attended moment and concurrent costs at unattended moments. Importantly, there are open questions regarding the timing and the precision of the temporal attentional window: (1) Do people use expectation to guide attention to a stimulus specific point in time or do they merely attend to its order in a sequence of events?; 2) Do the effects of temporal attention vary as a function of the precision with which people attend to the stimulus?

It has been proposed that expectations can guide temporal attention (review, Nobre & van Ede, 2018, 2023). Research suggests that temporal attention is directed towards a behaviorally relevant target when it is expected. When the target has not appeared at the expected moment, observers may shift their attention to the next possible time frame when an event may occur.

However, temporal attention and expectation have been used interchangeably or when defined as different mechanisms, they have not been independently manipulated (e.g., Capizzi et al., 2023; Chauvin et al., 2016; Correa et al., 2006; Coull, 2004; Coull & Nobre, 1998; Griffin et al., 2001). Thus, a more parsimonious explanation is that hazard rate–the updated expectation with increasing probability of an event occurring, given that it has not yet occurred–can account for these performance differences without necessarily invoking attention allocation. To investigate these alternative hypotheses, attention and expectation should be independently manipulated, to enable the comparison of the effects of attention at expected and unexpected time points.

Attention and expectation both improve performance but modulate neural responses in an opposite manner. In the visual system, temporal attention enhances neural responses to stimuli (Correa et al., 2006; Schroeder & Lakatos, 2009), whereas temporal expectations suppress these responses (Nara et al., 2021). In the auditory system, they interact at the neural level, expectation suppresses MEG beta-band oscillations for unattended- but not for attended-stimuli (Todorovic et al., 2015). Unfortunately, the potential perceptual interactions could not be examined because performance was at ceiling. In any case, in the visual system, it is unknown whether attention and expectation interact at the neural or behavioral level.

Here, we investigated the benefits of voluntary temporal attention on behavior while manipulating temporal expectation via precision and hazard rate (**Figure 1**). We implemented a protocol by combining temporal attention (Denison et al., 2017, 2021) and expectation (Todorovic et al., 2015; Todorovic & Auksztulewicz, 2021) manipulations. In our protocol, <u>temporal attention</u>–prioritization of a behaviorally relevant moment–is allocated based on the instructions to selectively process a target from two competing stimuli; temporal expectation–ability to guide behavior based on event probability–is manipulated via <u>temporal</u> <u>precision</u>, such that when event timing is certain precision is high, and <u>hazard rate</u> (used to index stimulus onsets earlier than, at or later than the expected moment). This protocol also enables us to distinguish whether people selectively attend to a stimulus at a specific point in time, or alternatively they merely rely on its order in a sequence, regardless of timing or expectations. Our results reveal that both temporal precision and hazard rate influence temporal attention, albeit in different manners.

**Figure 1.**
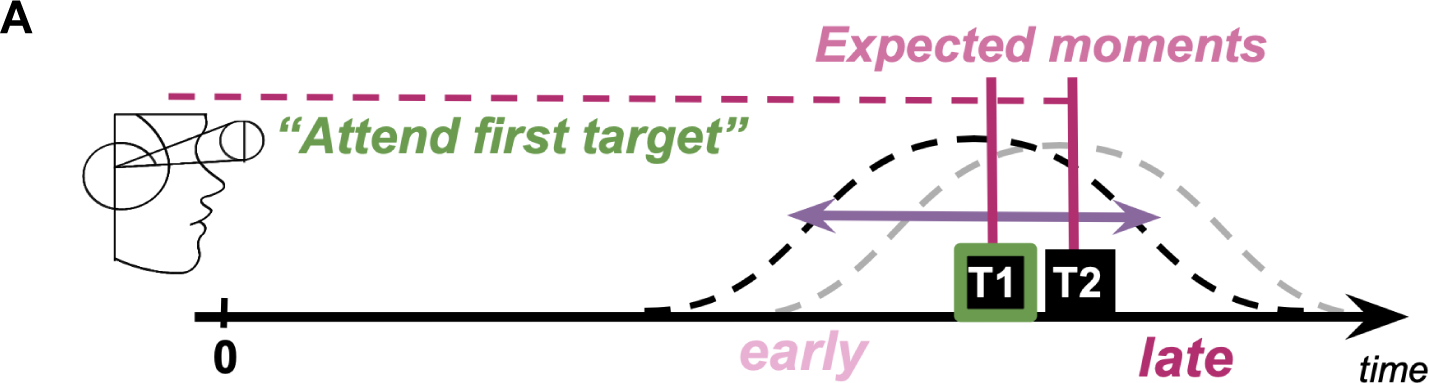
Illustration of the definitions and experimental manipulations of temporal attention, hazard rate and precision. The brain’s temporal processing capacity is limited, and it is a challenge to process the two brief sequential targets perfectly well. **Temporal Attention** is prioritization of a behaviorally relevant moment, and is allocated based on the instructions to selectively process a target that briefly appears (first target, T1, depicted here as an example), and ignore the subsequent one (T2). Given an event with a temporal probability distribution (distribution of possible visual event onsets), we can characterize time relative to the most expected time point: “early”, “expected”, and “late” time points. **Hazard Rate** characterizes the increase in probability of the onset as the time passes. **Temporal Precision** describes the inverse of the variability of temporal distribution.

## METHODS

### Observers

Sixteen observers (10 females, six males, aged 22-34 years), including one author have participated in the experiment. The number of observers needed was determined by a power analysis using G*Power software (Faul et al., 2007). The effect size was set to *ηG*2 = 0.14, which was reported in a previous paper with the same experimental protocol, and comparable number of manipulations (Fernández et al., 2019), and the sample size was evaluated at 80% power for the potential interaction between temporal attention and expectation.

All observers provided written informed consent and had normal or corrected-to-normal vision. All experimental procedures were in agreement with the Helsinki declaration and approved by the University Institutional Review Board.

### Apparatus

Stimuli were generated using an Apple iMac (3.06 GHz, Intel Core 2 Duo) and MATLAB 2012b (Mathworks, Natick, MA, USA) along with the Psychophysics Toolbox (Brainard, 1997; Kleiner et al., 2007), and presented on a CRT monitor that was color-calibrated (1280 × 960 screen resolution, 100-Hertz refresh rate). Observers were seated 57 cm from the display with their head movements limited by a chinrest. The Eyelink 1000 eye tracker (SR Research, Ottawa, Ontario, Canada) was employed to record eye position and perform online eye tracking to maintain central fixation throughout the trials. If a fixation break due to a blink occurred or if the eye position deviated more than 1° from the center of the screen between the ready cue and the response cue, the trial would be aborted and subsequently repeated at the end of each experimental block. Observers could blink or move eyes after the response cue and during the intertrial interval.

### Stimuli

Stimuli were displayed on a uniform medium gray background. A fixation circle (subtending 0.15° dva) was presented at the center of the screen. The placeholders were four small black circles (0.2°) placed at corners of an imaginary square (side length=2.2°) centered at the screen center.

Target stimuli were Gaussian-windowed (standard deviation of 0.3°) sinusoidal gratings (spatial frequency = 4 cpd) with random phase presented at full contrast. Each target was tilted clockwise (CW) or counterclockwise (CCW) from the vertical or horizontal axis. The tilt was titrated for each observer and for each target interval independently.

Auditory stimuli were presented through the speakers. The attentional precue was a 200-ms auditory tone, either a sinusoidal wave, or a complex waveform that is a combination of sinusoidal waves with frequencies ranging from 50-400 Hz. A high-frequency sinusoidal tone (800 Hz) indicated the first target, a low-frequency sinusoidal tone (440 Hz) indicated the second target, and the complex tone was uninformative regarding the target (neutral trials). The same tones (except for the neutral tone) were used as the response cue at the end of the trial to indicate the first (T1) or the second target (T2).

### Experimental Procedure

The experimental protocol was adapted from previous endogenous temporal attention literature (Denison et al., 2017, 2021; Fernández et al., 2019) and expectation literature (Todorovic et al., 2015). In a two alternative forced-choice task, observers were asked to report the orientation of one of the two Gabor stimuli. Throughout the experiment, the two targets were presented sequentially at the center of the screen, while the observers were fixating at that location.

Figure 2.B illustrates the experimental protocol. At the beginning of each trial, an auditory precue was presented to indicate whether to attend to the first target (T1), second target (T2) or both targets. Each target was presented for 30 ms with a stimulus onset asynchrony (SOA) of 250 ms. The onset of the targets relative to the precue was variable throughout the experiment, and was controlled throughout each session to manipulate temporal precision. An auditory 200-ms response cue was presented 500 ms after the second target onset. The observers were allowed to respond after the go cue, when the fixation brightness changed 800 ms after the response cue onset. In neutral trials, the response cue indicated first or the second Gabor with 50% probability, and in the other trials, precue was 100% valid and the response cue always matched the precue. Feedback was presented for 500 ms after each trial (A red minus after an incorrect response, or a green plus after a correct response). The observers had no time limit to respond, and were instructed to prioritize accuracy over speed when responding. The intertrial interval was jittered between 1700 and 1100 ms.

**Figure 2.**
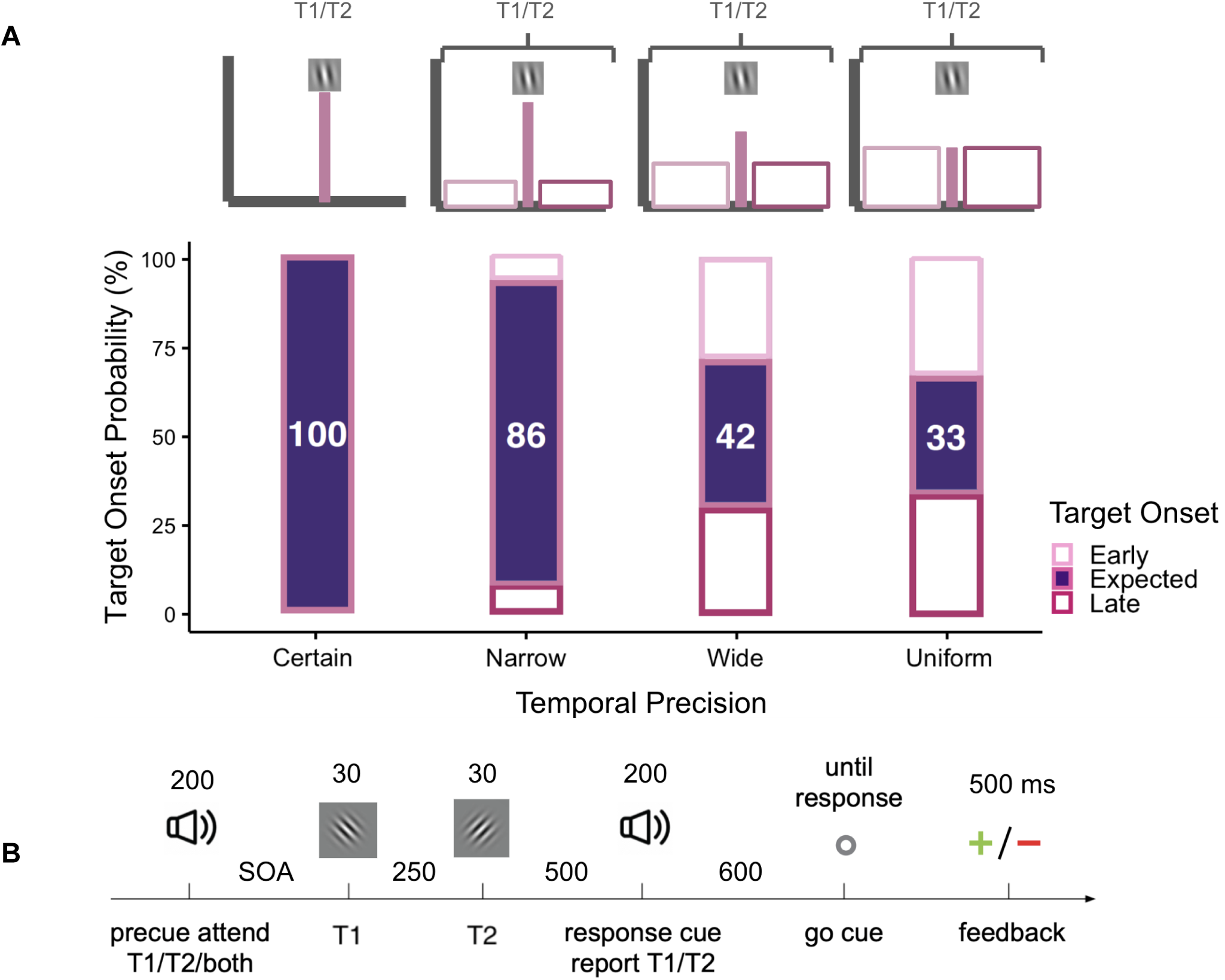
**(A)** Temporal precision of the stimulus onset was manipulated across sessions, and ranged from certain (no variability, highest precision) to uniform (lowest precision) (adapted from Todorovic et al., 2015)). Values inside the purple bars represent the percentage of trials that the stimuli appeared at the expected moment. For the certain condition, the targets appeared at the expected moment in 100% of the trials. With decreasing precision, targets could also appear earlier or later than this expected time point. Percentage of expected trials dropped with decreasing precision, 86% of the narrow, 42% for the wide and 33% for the uniform conditions. **(B)** Psychophysical procedure (adapted from Denison et al., 2017 to test visual performance at the attended and unattended time points when the stimulus onset is variable. Precue was either neutral, or indicated which target’s orientation will be asked to report at the end of the trial (response cue). Observers had unlimited time to respond, and received feedback based on the accuracy of the response.

Eye-tracking was used throughout the experiment, except during the practice session. Participants were instructed to avoid making eye movements and to focus on the center of the screen. If a participant looked away from the center of the screen or blinked during a trial, the trial was skipped and presented again at the end of the block. However, participants were free to make eye movements and blink after the response cue and before the pre-cue of the subsequent trial, defined as their response window.

Temporal precision within an experimental session was manipulated by changing the variability of the stimulus onsets relative to the precue, adapted from a previous neuroimaging study investigating the interaction between temporal expectation and temporal attention (Todorovic et al., 2015; Todorovic & Auksztulewicz, 2021). Figure 2.A illustrates the temporal distributions that created uncertainty–different levels of temporal precision. There were four temporal precision conditions in our design: certain, narrow, wide, and uniform. We tested visual performance at the same levels of stimulus probability as in the literature (42% and 86%), and added two additional temporal precision levels (33% and 100%), In all conditions, the probability distributions of T1 and T2 onsets were centered at 1400 and 1650 ms respectively after the precue onset (expected moments). And the probability of T1 appearing at 1400 ms, and T2 at 1650 ms after the precue onset was either 100% (certain), 86% (narrow), 42% (wide), and 33% (uniform, lowest precision) respectively, which determined the level of temporal precision. For each precision condition, we collected data for 1, 2, 3, or 4 sessions for certain, narrow, wide and uniform conditions respectively, to be able to have enough data points at the mean point (expected moment) of the temporal distribution. The temporal distributions were explained to the observers prior to each experimental session.

Except for the 100% certain condition, there was a time window (1200-1600 ms for T1, and 1450-1850 ms for T2) when the stimuli could occur, which enabled us to test the performance at earlier and later than the expected moment (midpoint of the temporal distributions, 1400 ms for T1 and 1650 ms for T2). In wide (42%) and uniform (33%) conditions, we had enough trials at the early and late time points that were comparable to the number of trials at the expected moment.

The order of the trials was randomized across the sessions, and the order of the sessions for temporal precision was randomized across participants. Each observer completed 7-10 sessions and 3568-4256 trials in total. Before each session, we titrated the neutral performance at the expected time points (1400 ms for T1 and 1650 ms for T2 after the precue) to be at 75% for both targets independently. Each experimental session started with the tilt threshold determined by the best PEST procedure (Lieberman & Pentland, 1982), and was adjusted on a block-by-block basis if the neutral accuracy significantly differed from the aimed baseline of 75% accuracy.

### Statistical analysis

Data analyses were performed with R (version 4.2.3; R Core Team, 2023), with ANOVA conducted using the ezANOVA package (Lawrence, 2016).

The discriminability index, d’ was computed by: z(hit rate) – z(false alarm rate). Correct discrimination of clockwise trials were categorized as hits and incorrect discrimination of counter-clockwise trials were categorized as false-alarms (Fernández & Carrasco, 2020; Zhang et al., 2019). We implemented a correction to avoid infinite values when computing *d*′, and added 0.5 to the number of hits, misses, correct rejections, and false alarms before computing *d*′ (Brown & White, 2005; Hautus, 1995).

We use median values for RTs (from the go-cue) throughout the analysis as the participants had infinite time to respond, causing the RT distribution to be skewed, which makes the median a less biased estimator for RT.

To capture the effects on accuracy and the speed of responses with a single metric, we also calculated the Balanced Integration Scores (BIS):

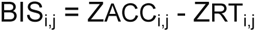

Here, BIS_i,j_ represents the difference between standardized mean correct accuracies and median RTs, affording equal importance to each. The subscripts, i and j, correspond to participant i’s performance for condition j (Liesefeld & Janczyk, 2019).

To provide a better summary of the attentional modulations across precision and expectation, we computed a benefit index using the BIS scores (adapted from Denison et al., 2017):

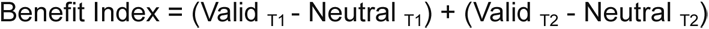

## RESULTS

### Temporal precision

To investigate whether temporal attention interacts with temporal precision, we analyzed the data where the stimuli occurred at the expected time points (1400 ms for T1 and 1650 ms for T2 relative to the precue onset). In those trials, the trial timings and experimental manipulations were exactly the same, except that those trials were embedded in sessions with different temporal probabilities (Certain, Narrow, Wide, Uniform conditions), resulting in different levels of temporal precision.

We performed three-way ANOVAs (2 target x 2 precue x 1 precision) on sensitivity (d’), criterion, reaction time (RT), and Balanced Integration Scores (BIS). The factors analyzed were target (T1, T2), cue validity for temporal attention (Valid, Neutral), and temporal distribution for the temporal precision (Certain, Narrow, Wide, Uniform; in which the target occurred at the expected moments with a respective probability of 100%, 86%, 42%, and 33%). We registered temporal distribution as a numeric variable in order to account for the spacing between the probability values (MacCallum et al., 2002).

Significant main effects for d’ were found for cue validity, F(1,15) = 5.180, p = 0.038, *η^2^*G = 0.029, and temporal distribution (Figure 3.A), F(1,15) = 9.341, p = 0.008, *η^2^*G = 0.080. Furthermore, a significant interaction between target and cue validity was found, F(1,15) = 5.757, p = 0.030, *η^2^*G = 0.031 (Figure 4.A). Holm-corrected pairwise t-tests showed that d’ was higher than Valid than Neutral condition for T1, p = 0.000, and there was no significant difference between those conditions for T2 (p = 0.520). Pairwise t-tests for temporal distribution also showed that d’ for Certain was higher than d’ for Wide, p = 0.031, and that d’ for Certain was higher than d’ for Uniform, p = 0.021. Overall, accuracy increased with temporal attention for the first target, and across targets, it gradually increased with temporal precision.

**Figure 3.**
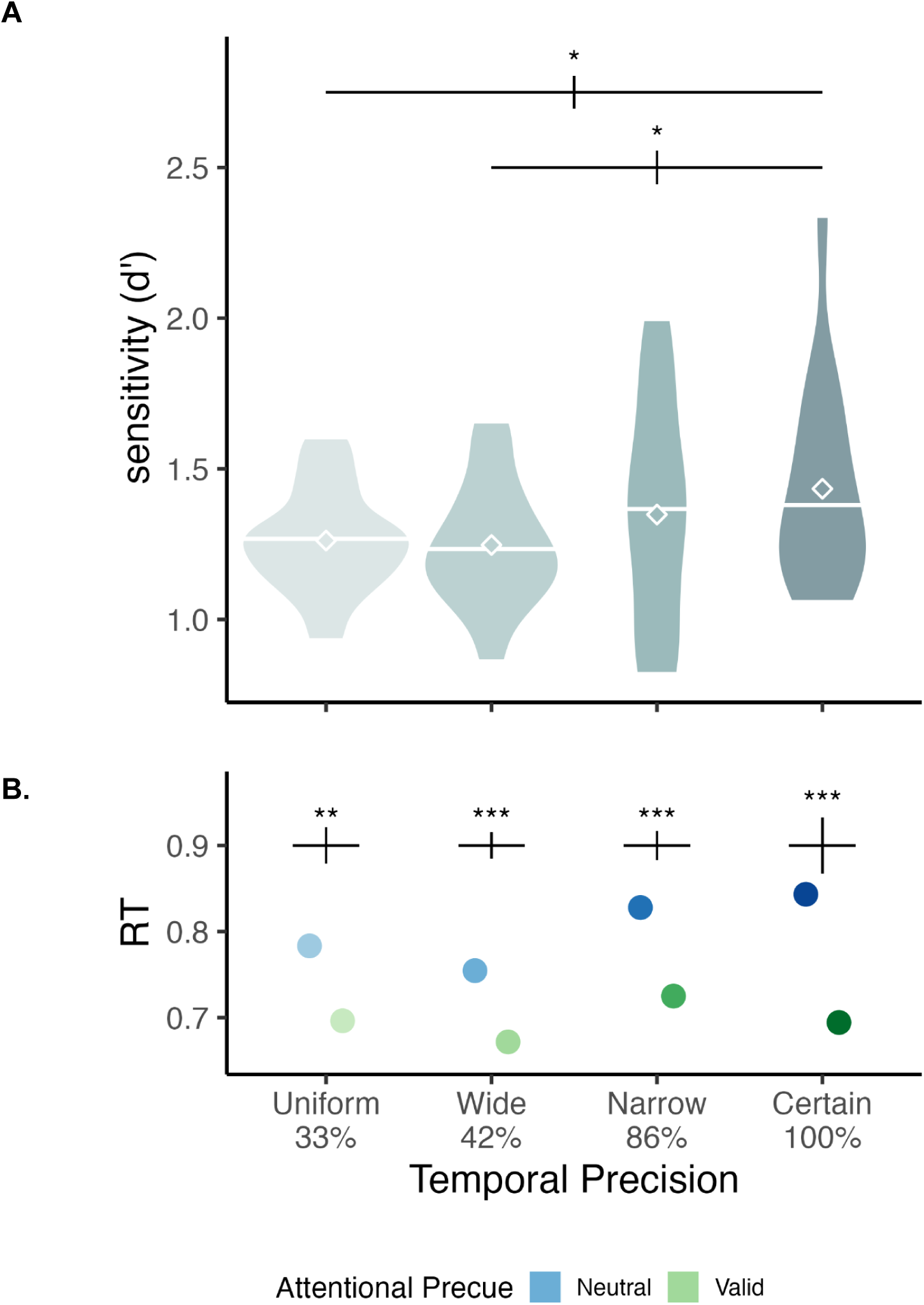
d’ and reaction time computed at the expected time point of all precision conditions. Only trials where T1 and T2 appeared at the expected time point (1400 and 1650, respectively) were included in the analysis. Error bars above each group of bars denote within-subject error rate with Morey correction (Cousineau, 2005; Morey, 2008). (A) Main effect of temporal precision on discriminability (d’). The results suggest a gradual increase in overall performance with increasing precision. (B) The interaction between cue validity and temporal precision on reaction time. The benefit of temporal attention was present across the levels, although it gradually increased with precision.

**Figure 4.**
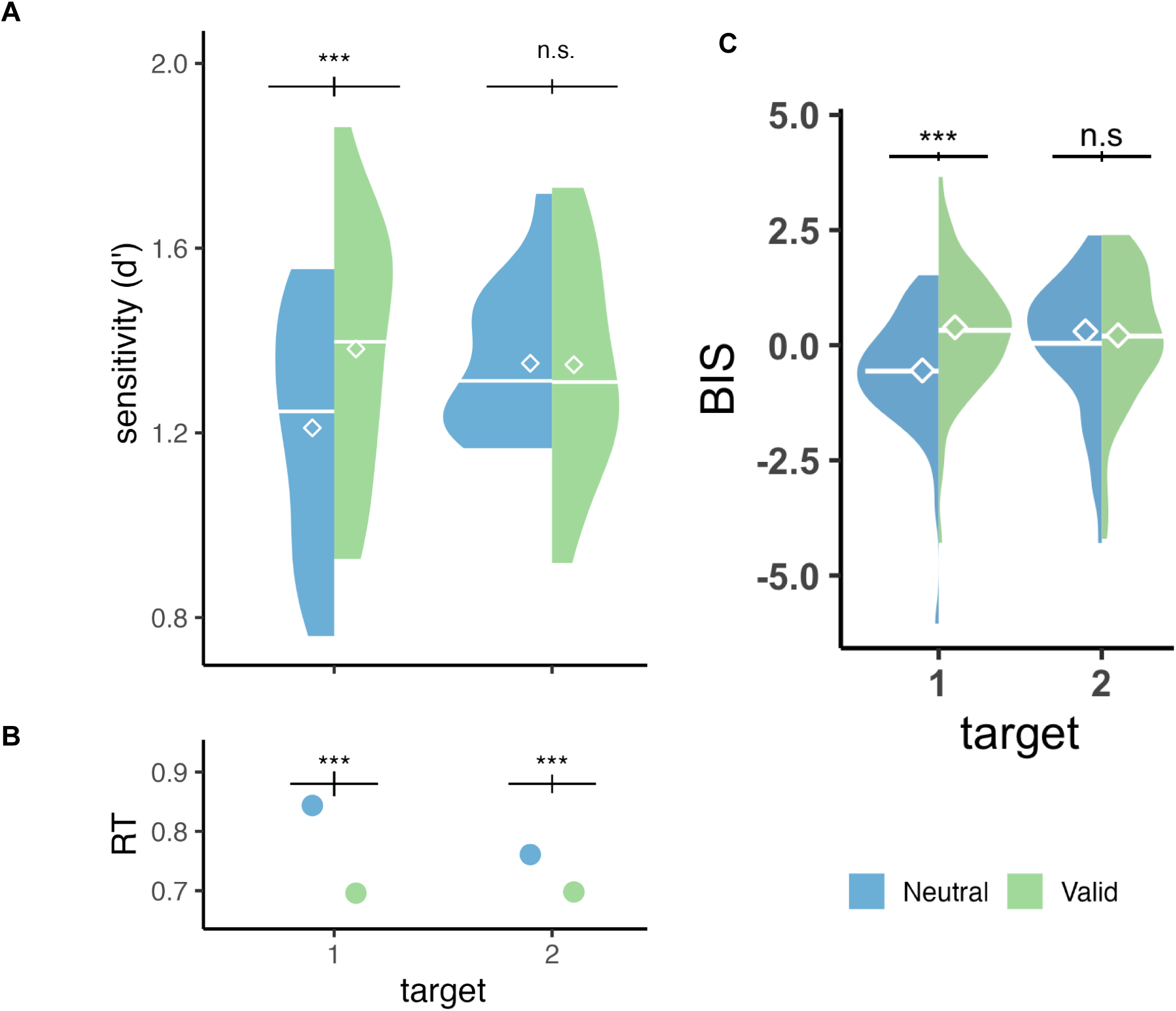
Performance (d, RT, BIS)’ computed at the expected moment of each temporal distribution (T1 onset 1400 ms, and T2 onset 1650 ms). Significant interaction between cue validity and target was present for d’, RT and BIS.

The ANOVA on the criterion revealed no significant main effects or interactions (All ps>0.1).

For RT, we found a significant main effect for cue validity, F(1,15) = 17.696, p = 0.001, *η^2^*G = 0.231. Additionally, significant interactions were found between cue validity and temporal distribution, F(1,15) = 5.869, p = 0.029, *η^2^*G = 0.014 (Figure 3.B), and between target and cue validity, F(1,15) = 36.979, p = 0.000, *η^2^*G = 0.046 (Figure 4.B). The holm-corrected pairwise comparisons revealed slower RTs in valid than neutral conditions (p = 0.000), and although this was present for both targets, it was more pronounced for T1 than T2 (both ps = 0.000). And the same effect, reduction in RT with attention was observed in all temporal precision conditions, even though it was the smallest under lowest precision (p = 0.001 for uniform (33%), p = 0.000 for other levels). In sum, temporal attention facilitated reaction times, and more so for the first target, and this attentional modulation increased with temporal precision.

For BIS, a significant main effect was found for cue validity, F(1,15) = 16.857, p = 0.001, *η^2^*G = 0.137. Moreover, significant interactions were found between target and cue validity, F(1,15) = 23.724, p = 0.000, *η^2^*G = 0.072 (Figure 4.C), between target and precision, F(1,15) = 5.134, p = 0.039, *η^2^*G = 0.039 (Figure 5.A), and between cue validity and precision, F(1,15) = 6.095, p = 0.026, *η^2^*G = 0.017 (Figure 5.B). Holm-corrected pairwise t-tests between cue validity and target revealed a higher BIS score for the T1 Valid than the Neutral condition, p = 0.000, and no effect for the T2, p = 0.140. Pairwise t-tests for cue validity and temporal distribution showed a significantly higher BIS score for the Certain Valid than the Neutral, p = 0.000, for the Narrow Valid than the Neutral condition, p = 0.004, and a statistical trend for the Wide Valid than the Neutral, whereas no effect was found for the uniform condition. Pairwise comparisons revealed that for T1, observers had a significantly higher performance with certain timing (100%) than uniform (p = 0.004) and wide temporal distributions (p = 0.013). For T2, there was no significant difference across the levels of temporal precision. To summarize, visual performance was improved by temporal attention and temporal precision for the first target. Moreover, across the targets, temporal attention and precision interacted, and the benefits of temporal attention increased with temporal precision.

**Figure 5.**
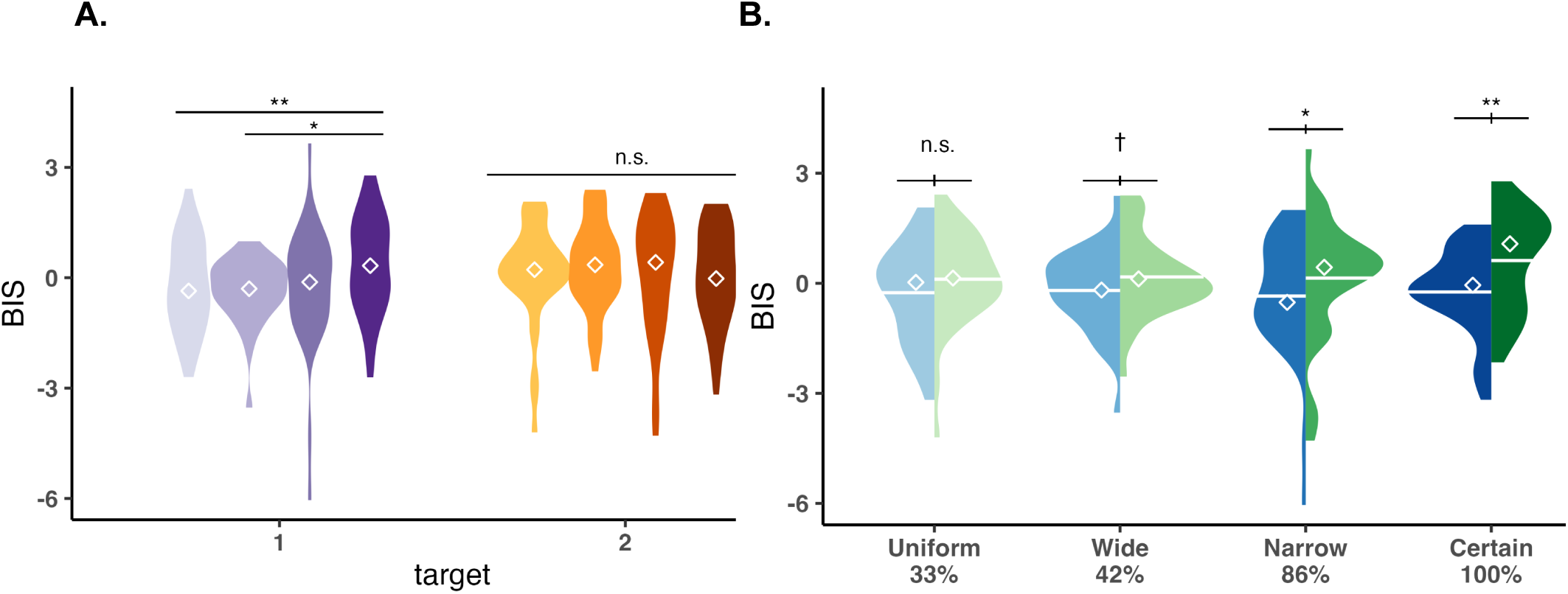
Temporal precision is indicated with the darkness of the colors. **A.** Significant interaction between target and temporal precision. The level of temporal precision affected T1, but not T2. **B.** Significant interaction between temporal precision and cue validity. Temporal attention benefits performance when target timing is predictable, and the magnitude of benefit increases with precision.

One way ANOVA on the benefit index revealed a significant effect of temporal precision, F(1,15) = 6.095, p = 0.026, *η^2^*G = 0.289 (Figure 6), and the pairwise comparisons showed a significant difference between the certain and uniform conditions (p=0.05). Moreover, t-tests comparing data to 0 (no benefit) revealed no significant effect for the uniform and wide temporal distributions (p>0.1), whereas significant effects on attentional benefits were found for narrow (p = 0.009) and certain (p=0.004) conditions. In sum, temporal attention benefits gradually increased with temporal precision, and were minimal under low temporal precision.

**Figure 6.**
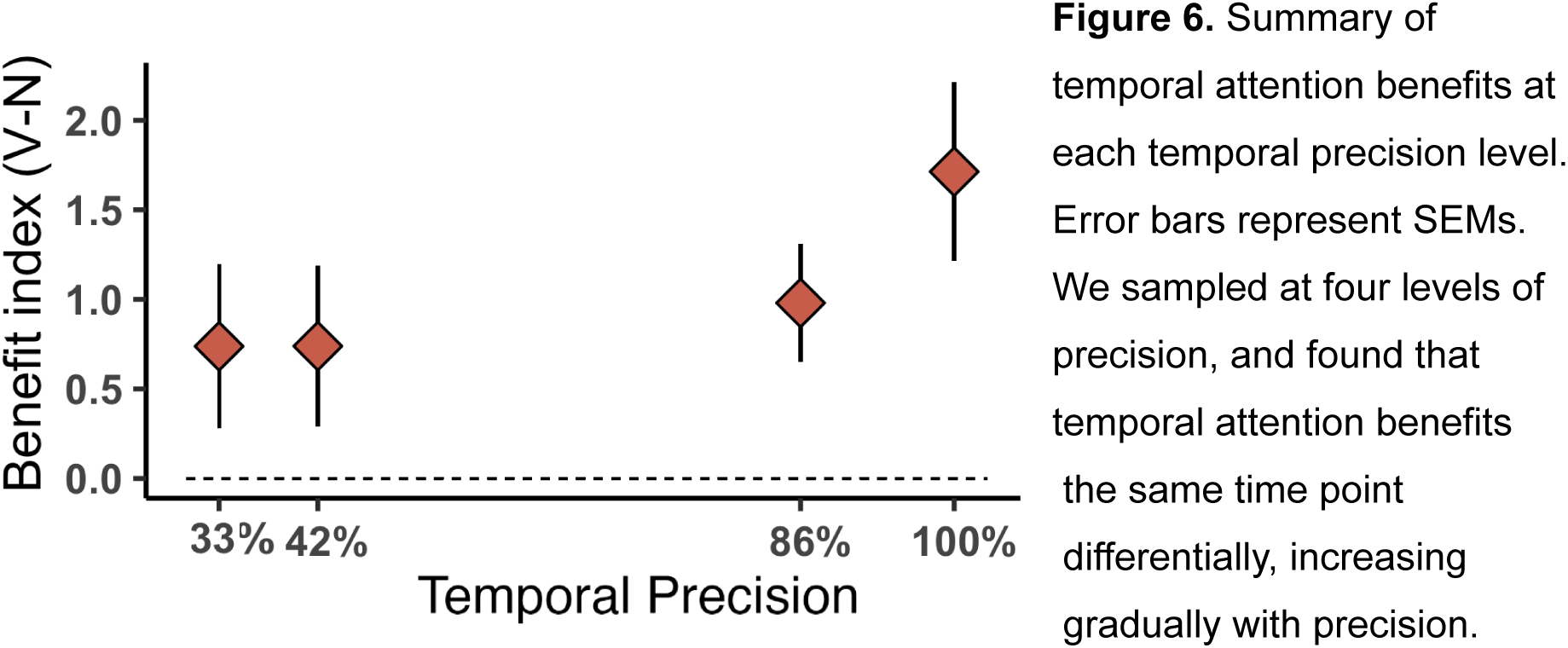
Summary of temporal attention benefits at each temporal precision level. Error bars represent SEMs. We sampled at four levels of precision, and found that temporal attention benefits the same time point differentially, increasing gradually with precision.

### Hazard rate

To analyze performance at expected and unexpected moments, we analyzed the entire time window in the wide (42%) and uniform (33%) conditions, as these two conditions had enough trials at the unexpected time points. We collapsed the data across these two conditions, as no significant difference was found between them in the previous step. We categorized the stimulus onsets relative to the expected moment: Early, Expected, Late onsets.

The potential interaction between temporal expectation and attention on d’, criterion, reaction time (RT), and balanced integration scores (BIS) was assessed by three-way ANOVAs (2×2×3). The factors were target (T1, T2), cue validity (Valid, Neutral), and the stimulus onset (Early, Expected, Late). For this analysis, we did not register stimulus onset as numerical, as the independent variables were evenly spaced (but results were the same when we did so).

The analysis on d’ revealed significant interactions between target and cue validity, F(1,15) = 9.705, p = 0.007, *η^2^*G = 0.021, and between cue validity and stimulus onset, F(2,30) = 4.656, p = 0.017, *η^2^*G = 0.011. A significant three-way interaction between target, cue validity, and stimulus onset was also present, F(2,30) = 3.537, p = 0.042, *η^2^*G = 0.015 (Figure 7.A). No significant main effect for the stimulus onset (p>0.1). A statistical trend for target was found (p=0.076). Holm-corrected pairwise t-tests for the three-way interaction revealed a significant difference between Neutral and Valid conditions for T1 at early onsets (p = 0.000), followed by a smaller effect at the expected onset (p = 0.034), and there was no effect of temporal attention at late onsets (p = 0.52). No significant effect of cue validity was found for T2 across the time window. Temporal attention improved discriminability for the first target, specifically when the target appeared earlier than, or at the expected moment, but not later.

**Figure 7.**
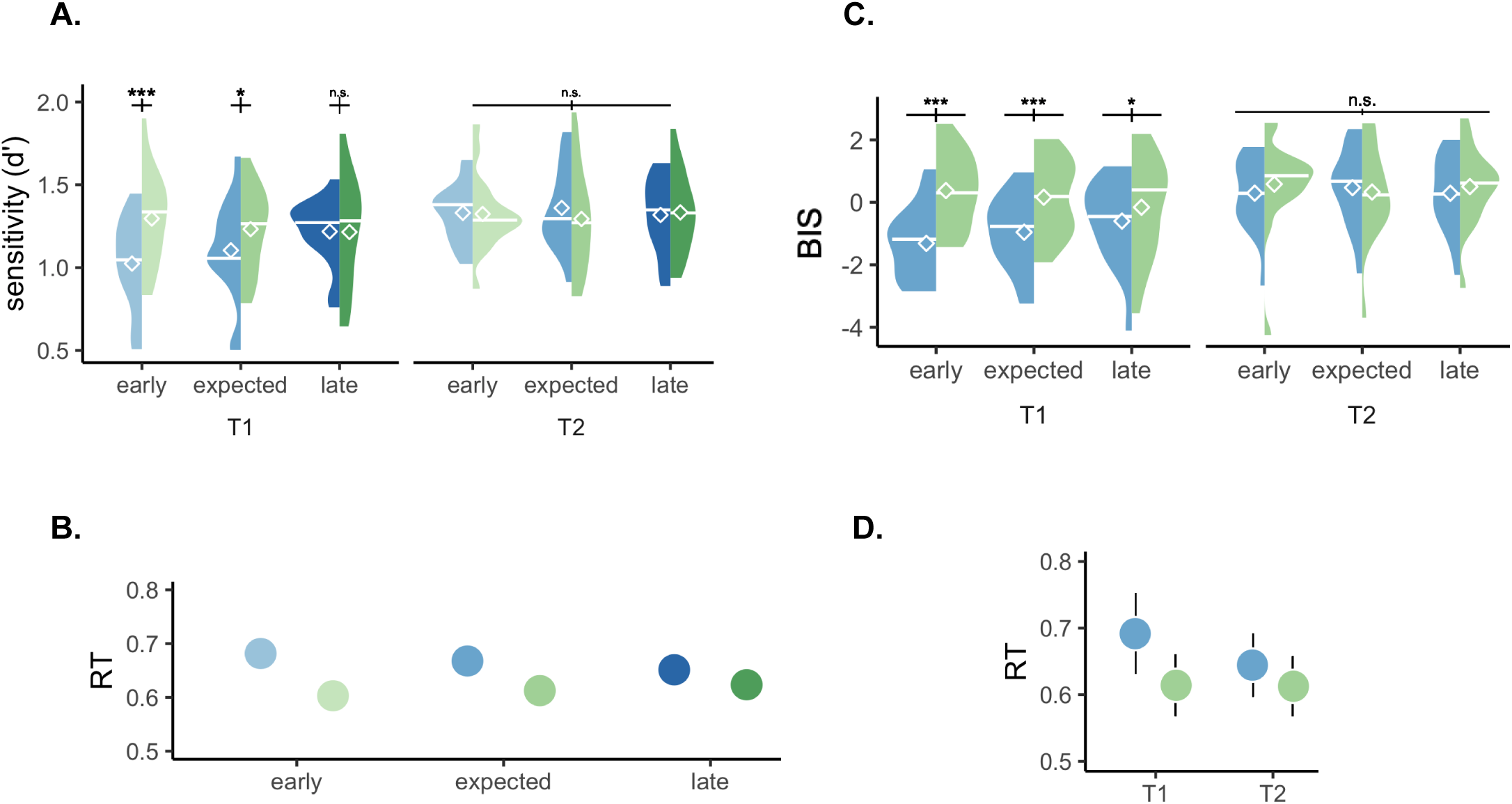
Performance was analyzed across the time window that the stimuli could appear (wide 42% and uniform 33% temporal precisions). **A & B.** Analyses on d’ and BIS revealed three-way significant interactions between target, stimulus onset and cue validity. Temporal attention improved performance for T1, but not T2. The improvements for T1 were absent after the expected moment. **C.** Benefits on RT were present across the time window (all ps<0.001), although this effect got smaller as the stimulus onset was delayed (revealed by an interaction between the stimulus onset and cue validity). **D.** Temporal attention benefits on RT were present for both targets (both ps<0.001), although was stronger for T1 than it was for T2, indicated by a significant interaction between target and cue validity.

A three-way ANOVA on the criterion did not reveal any significant main effect or interaction (All ps>0.1).

With regard to RT, a significant main effect was observed for cue validity, F(1,15) = 17.670, p = 0.0007, *η^2^*G = 0.041, and for target F(1,15) = 6.969, p = 0.018, *η^2^*G = 0.025. Significant interactions were found between target and cue validity, F(1,15) = 16.195, p = 0.001, *η^2^*G = 0.008, and between cue validity and stimulus onset, F(2,30) = 7.090, p = 0.003, *η^2^*G = 0.0008 (Figure 7.B). Pairwise t-tests between stimulus onset, cue validity, and target revealed lower RT in valid than neutral conditions for both T1 and T2 across the entire temporal window (all ps<0.001), but this effect was stronger for T1 than T2, and stronger at early, than expected, followed by the late onsets. Temporal attention facilitated RT, and this facilitation was the strongest for the early targets for T1.

For BIS, significant main effects were found for target, F(1,15) = 7.803, p = 0.014, *η^2^*G = 0.093, and cue validity, F(1,15) = 7.351, p = 0.016, *η^2^*G = 0.051. There were also significant interactions between target and cue validity, F(1,15) = 26.289, p = 0.000, *η^2^*G = 0.034 (Figure 7.D), and between cue validity and stimulus onset, F(2,30) = 8.085, p = 0.002, *η^2^*G = 0.012, and a three-way interaction between target, cue validity, and stimulus onset, F(2,30) = 4.546, p = 0.018, *η^2^*G = 0.01 (Figure 7.C). Pairwise comparison between cue validity for the target at each time point showed a higher BIS score for the T1 Valid than Neutral condition that got smaller with stimulus onset, with the strongest difference at early onsets (p = 0.000), followed by the expected moment (p = 0.000), and followed by a statistical trend at late onsets (p = 0.085). No significant difference was found for T2 across the time window (all ps>0.1). Overall, we found that temporal attention improved performance for T1, the strongest in the early time window, followed by the expected moment, and then with a minimal improvement when targets appeared later than expected.

## DISCUSSION

This study reveals an interaction between voluntary temporal attention and temporal expectation. The benefits of temporal attention on performance depended on expectation: They gradually increased as a function of temporal precision; conversely, and surprisingly given previous findings, they decreased with the onset delay of the behaviorally relevant stimulus, despite its increasing probability of appearance.

### People use expectation suboptimally to guide attention to a stimulus specific point in time

Relevant moments can be selected and prioritized through endogenous temporal attention when events appear at certain time points, resulting in perceptual benefits and costs at attended and unattended times, respectively (Denison et al., 2017, 2021; Fernández et al., 2019). The protocols in these studies isolated temporal attention and expectation by keeping the timing of the sequential stimuli constant. Thus, whether expectation needs to be temporarily precise, or is even necessary for temporal attention allocation was an open question. Perceptual tradeoffs in these experiments could have been attained merely by selecting a visual event based on the order in which they appear in a sequence, regardless of temporal expectation (e.g., the first event among the sequence), or by considering temporal expectation, by estimating the probabilities of when events would occur, and directing attention to a specific time point beforehand (e.g., 1 second after the precue). The protocol we used here enabled us to disentangle these alternatives. According to the former possibility, temporal attention would be sustained and the results would reflect uniform benefits across the tested time window. According to the latter possibility, temporal attention should peak at the expected moment, and would gradually increase as a function of precision of the expectation. Our results revealed that temporal precision modulated attentional benefits, hence expectation needs to be temporally precise. Moreover, temporal attention benefits were not constant over time across the hazard function, indicating that attentional allocation goes beyond the stimulus order in the sequence. Interestingly, temporal attention benefit was higher for earlier than the expected time points and gradually decreased with onset delay. These results indicate that temporal attention cannot be sustained across time despite constant competition between two sequential stimuli, and suggest that it should be precisely allocated.

We interpret the higher benefit of temporal attention before the expected time to reflect suboptimal allocation. Optimal allocation of temporal attention would have yielded the highest benefits at the expected moment, and symmetrical benefits at earlier and later moments. Instead, we found an *attentional undershoot–*the highest benefits at early moments, followed by the expected moment, and minimal benefit at the later window. In the current design the late stimulus onset could not be predicted at the beginning of the trial, and if attentional resources were spent they could not be reallocated. In contrast, temporal attention can benefit performance at the same late time window when there is no temporal uncertainty and resources could be saved to be deployed at the appropriate time (Denison et al., 2017).

### Temporal attention benefits vary as a function of temporal precision

The dynamic normalization model of temporal attention explains the temporal attention effects when stimulus timing is certain, and assumes precise selection of those behaviorally time points (Denison et al., 2021). It has been pointed out that this model does not differentiate whether the behavioral modulations are attained through differential anticipation before the two stimuli, and/or differential prioritization and filtering of the relevant stimulus during (or after) sensory encoding (Shalev & van Ede, 2021). Our new protocol and findings address this point. They reveal that temporal attention is not sustained across time, and a gradual decrease in benefits with a delayed stimulus onset suggests that temporal attentional selection occurs prior to the stimulus onset. We also highlight that a read out effect after the sensory encoding would result in the same performance for valid and neutral conditions in the prior (Denison et al., 2017, 2019; Fernandez et al., 2019) and the present studies.

Temporal expectation has been shown to have different sources in the auditory domain; hazard rate evokes earlier responses in early auditory regions and superior temporal gyri, whereas temporal precision-evoked responses have a delayed onset in the parietal cortex (Todorovic & Auksztulevicz, 2021). In the current study, we use a similar protocol to manipulate expectations, so that they increase as a function of both temporal precision and hazard rate. We found that the benefits of temporal attention interact with expectation, but the direction of the modulation is opposite for the two sources. On the one hand, with increasing expectations, based on temporal precision, attentional benefits gradually increase. On the other hand, temporal attention benefits decrease as the stimulus onsets are delayed via the hazard rate. These behavioral findings provide further evidence for the hypothesized distinction between the two sources of temporal expectation.

Previous studies concluding that expectations can guide temporal attention report a temporal asymmetry when there is a mismatch between the participants’ expectation and the target onset; lower performance when a target appears earlier than at an expected moment than when it appears later than it. The authors interpret the findings as expectation guidance of attention to an early time point in the trial, and when the cue is invalid, hazard rate guides observers to voluntarily reorient attentional resources to a later time point (Capizzi et al., 2023; Correa et al., 2006; Coull & Nobre, 1998; Nobre & van Ede, 2018, 2023). However, these two processes are not independently manipulated in their experiments. Our study distinguishes between temporal attention and expectation, as between two sources of expectation: We found that expectations that are known beforehand–temporal precision–do indeed modulate temporal attention. Our results revealed that temporal attention cannot be reoriented with evolving probabilities when the target onset is delayed. Had temporal attention been flexibly reoriented guided by the hazard rate, we would have observed temporal attention benefits at the late time window. In contrast, our results suggest that previous behavioral improvements could be explained by temporal expectations, without invoking the allocation of temporal attention. We note that the lack of improvements at late intervals in our design could be due to the shorter temporal window (400 ms) than in the designs of the aforementioned studies (500-1000ms). But how much time is needed for attention recovery is an open question.

Exogenous temporal attention–involuntary selection of specific time points following brief salient changes–interacts with expectation. It improves performance when the stimuli are presented at a later time point, but not earlier, in the trial when stimuli onset probability is uniform (Duyar et al., 2023). The current findings show that endogenous, voluntary temporal attention interacts with expectations in an opposite manner; they were pronounced at the early window and minimal at the late window. These opposite findings indicate a distinction between exogenous and endogenous attention, akin to the distinction of these subsystems in spatial attention (e.g., Fernández et al., 2022, 2023; Jigo et al., 2021).

In conclusion, this study highlights that temporal attention and expectation are different mechanisms whose effects depend on the contextual and goal demands. Temporal expectation can guide attention; the more precise the timing the higher the benefits; however, attention is suboptimally allocated at earlier than expected moments, and once allocated, it cannot immediately be reoriented to a later time point if visual events are delayed.

## OPEN PRACTICES STATEMENT

The data and codes for the study is available upon request from the authors.

